# Amyloidogenic Proteins Drive Hepatic Proteostasis Remodeling in an Induced Pluripotent Stem Cell Model of Systemic Amyloid Disease

**DOI:** 10.1101/358515

**Authors:** Richard M. Giadone, Derek C. Liberti, Taylor M. Matte, Jessica D. Rosarda, Celia Torres-Arancivia, Sabrina Ghosh, Jolene K. Diedrich, Sandra Pankow, Nicholas Skvir, J.C. Jean, John R. Yates, Andrew A. Wilson, Lawreen H. Connors, Darrell N. Kotton, R. Luke Wiseman, George J. Murphy

**Affiliations:** Center for Regenerative Medicine of Boston University and Boston Medical Center, Boston, MA, USA; Alan and Sandra Gerry Amyloid Research Laboratory, Amyloidosis Center, Boston University School of Medicine, Boston, MA, USA; Department of Molecular Medicine, The Scripps Research Institute, La Jolla, CA, USA; The Pulmonary Center and Department of Medicine, Boston University School of Medicine, Boston, MA, USA; Section of Hematology and Oncology, Department of Medicine, Boston University School of Medicine, Boston, MA, USA

**Keywords:** hereditary amyloidosis, pluripotent stem cells, gene editing, single cell transcriptomics, hepatic disease

## Abstract

Systemic amyloidosis represents a class of disorders in which misfolded proteins are secreted by effector organs and deposited as proteotoxic aggregates at downstream target tissues. Despite being well-described clinically, the contribution of effector organs such as the liver to the pathogenesis of these diseases is poorly understood. Here, we utilize a patient-specific induced pluripotent stem cell (iPSC)-based model of hereditary transthyretin (TTR) amyloid disease (ATTR amyloidosis) in order to define the contributions of hepatic cells to the distal proteotoxicity of secreted TTR. To this end, we employ a gene correction strategy to generate isogenic, ATTR amyloidosis patient-specific iPSCs expressing either amyloidogenic or wild-type TTR. We further utilize this gene editing strategy in combination with single cell RNAseq to identify multiple hepatic proteostasis factors, including many components of adaptive unfolded protein response (UPR) signaling pathways, whose expression correlates with the production of destabilized TTR variants in iPSC-derived hepatic cells. We further demonstrate that enhancing ER proteostasis within ATTR amyloidosis iPSC-derived hepatic lineages via stress-independent activation of aforementioned adaptive UPR signaling preferentially reduces the secretion of destabilized amyloidogenic TTR. Together, these results suggest the potential of the liver to chaperone-at-a-distance and impact pathogenesis at downstream target cells in the context of systemic amyloid disease, and further highlight the promise of UPR modulating therapeutics for the treatment of TTR-mediated and other amyloid diseases.

## INTRODUCTION

Systemic amyloid disease represents a class of devastating protein folding disorders affecting over 1 million individuals worldwide (1–5). In these diseases, proteins containing destabilizing mutations are produced and secreted from an effector organ into circulation. In the blood, these proteins undergo misfolding and subsequent aggregation into toxic oligomers and amyloid fibrils, which deposit at distal target organs resulting in cellular death and organ dysfunction. Systemic amyloid diseases can result from the pathogenic misfolding of over 15 structurally distinct proteins, a majority of which are synthesized by the liver. A prominent example of this family of diseases is hereditary transthyretin amyloidosis (ATTR amyloidosis).

Hereditary ATTR amyloidosis is a complex autosomal dominant disorder that can result from over 100 possible mutations in the transthyretin (*TTR*) gene (6–9). In normal conditions, TTR is produced chiefly by the liver, where it forms a homotetramer and is then secreted, allowing for the transport of thyroxine and retinol binding protein throughout circulation (6–9). In patients with ATTR amyloidosis, TTR mutations decrease the stability of the tetramer and result in monomerization and subsequent misfolding of TTR variants. Amyloid-prone monomers then aggregate to form proteotoxic low-molecular weight oligomers and eventually hallmark congophilic amyloid fibrils at downstream target tissues including the heart and peripheral nerves (6–9). Current standards of care for patients with the disease include small molecule kinetic stabilizers tafamidis and diflunisal, which work to stabilize the tetrameric protein and limit monomerization and downstream fibril formation (10–14). Despite some success in clinical trials for both compounds and FDA approval for the use of tafamidis in the treatment of TTR-related cardiomyopathy, not all patients respond equally and effectively to these medicines. This can be attributed to the inherited deleterious *TTR* mutation as well as the underlying genetic background of the individual (10–15). Due to their multi-tissue etiologies and age-related trajectories, systemic amyloid diseases like ATTR amyloidosis have proven difficult to study in a physiologically-relevant way. At the same time, no mouse model currently recapitulates key aspects of human TTR amyloid pathology (16–20). To better understand disease pathogenesis in the genetic context of affected patients, we leveraged our previously described, patient-specific induced pluripotent stem cell (iPSC)-based model for studying the disease. In this system, TTR amyloid disease-specific iPSCs are differentiated into effector hepatocyte-like cells (HLCs) that produce and secrete destabilized, amyloidogenic mutant TTR that can then be used to dose a myriad of iPSC-derived, disease-associated target cells (18–20).

Traditionally, the livers of ATTR amyloidosis patients have been thought to be unaffected throughout disease pathogenesis, as toxicity occurs at downstream target organs such as the heart and peripheral nerves (1–9). Despite this however, recent studies suggest the capacity for the liver to contribute to the deposition of amyloidogenic proteins at distal target tissues. In line with this, recipients of domino liver transplantations (DLTs), where an individual in end-stage liver failure receives a liver from an ATTR amyloidosis patient, show accelerated TTR amyloid disease pathogenesis, wherein TTR fibrils accumulate on target organs in fewer than 10 years (21–26). Furthermore, *in vivo* mouse experiments have shown that the deposition of TTR on the hearts of old mice correlates with altered expression of numerous genes in the liver associated with the regulation of hepatic proteostasis (27). These results implicate the liver in the pathogenesis of systemic amyloid diseases such as ATTR amyloidosis. *Despite these observations however, the molecular and cellular changes in the liver that contribute to the toxic aggregation of misfolded TTR in distal tissues remains unclear*.

Interestingly, significant evidence highlights an important role for endoplasmic reticulum (ER) proteostasis in regulating the secretion and subsequent aggregation of amyloidogenic proteins such as TTR (28–32). The capacity of the ER proteostasis environment to secrete destabilized, aggregation-prone TTR variants has been directly implicated in the onset and pathogenesis of ATTR amyloidosis-related disorders. Furthermore, activation of adaptive IRE1/XBP1s or ATF6 arms of the unfolded protein response (UPR) – the predominant signaling pathway responsible for regulating ER proteostasis – have been shown to reduce the secretion and subsequent aggregation of structurally diverse amyloidogenic proteins (including TTR) (29, 31). Despite this, the physiological importance of ER proteostasis and the UPR in ATTR amyloidosis remains poorly defined.

Here, we utilize our laboratory’s previously described patient-specific iPSC-based model of ATTR amyloidosis to investigate the contribution of proteostasis and hepatic disease modifying factors to the distal toxicity observed in patients with the disease. By utilizing gene editing in combination with single cell RNA sequencing (scRNAseq), we define distinct transcriptional profiles in syngeneic corrected and uncorrected ATTR amyloidosis iPSC-derived HLCs that correlate with expression of a destabilized, amyloidogenic TTR variant. Through these efforts, we show that expression of the most proteotoxic, destabilized, disease-associated TTR mutant in HLCs increases expression of genes and pathways inversely implicated in the toxic extracellular aggregation of TTR, including transferrin and UPR target genes. To assess the consequence of functional activation of adaptive UPR signaling within hepatic cells expressing mutant TTR, we generated an ATF6-inducible patient-specific iPSC line. We further utilize this tool to demonstrate that exogenous ATF6 activation preferentially reduces the secretion of mutant, amyloidogenic TTR relative to the wild-type protein.

Herein, we demonstrate that hepatic expression of amyloidogenic TTR results in differential expression of proteostasis factors known to protect the extracellular space from toxic protein aggregation.

## RESULTS

### TTR is a top differentially expressed gene throughout human hepatic specification

Recent work from our group demonstrated the emergence of a stage-dependent disease signature of hepatic-specified pluripotent stem cells (PSCs) (33). In these experiments, microarray analyses were performed on cells isolated at days 0, 5, and 24 of hepatic differentiation. Post-hoc analysis of this data revealed that *TTR* was the second most differentially expressed gene comparing differentiated HLCs to PSCs (**Fig. 1A**). To confirm this, qRT-PCR was performed on RNA isolated from day 24 HLCs, demonstrating significant upregulation of *TTR* as compared to undifferentiated iPSCs (**Fig. 1B**). These data demonstrate that TTR is a robust marker of hepatic specification and can be used to normalize stem cell-derived hepatic differentiations.

**FIGURE 1.**
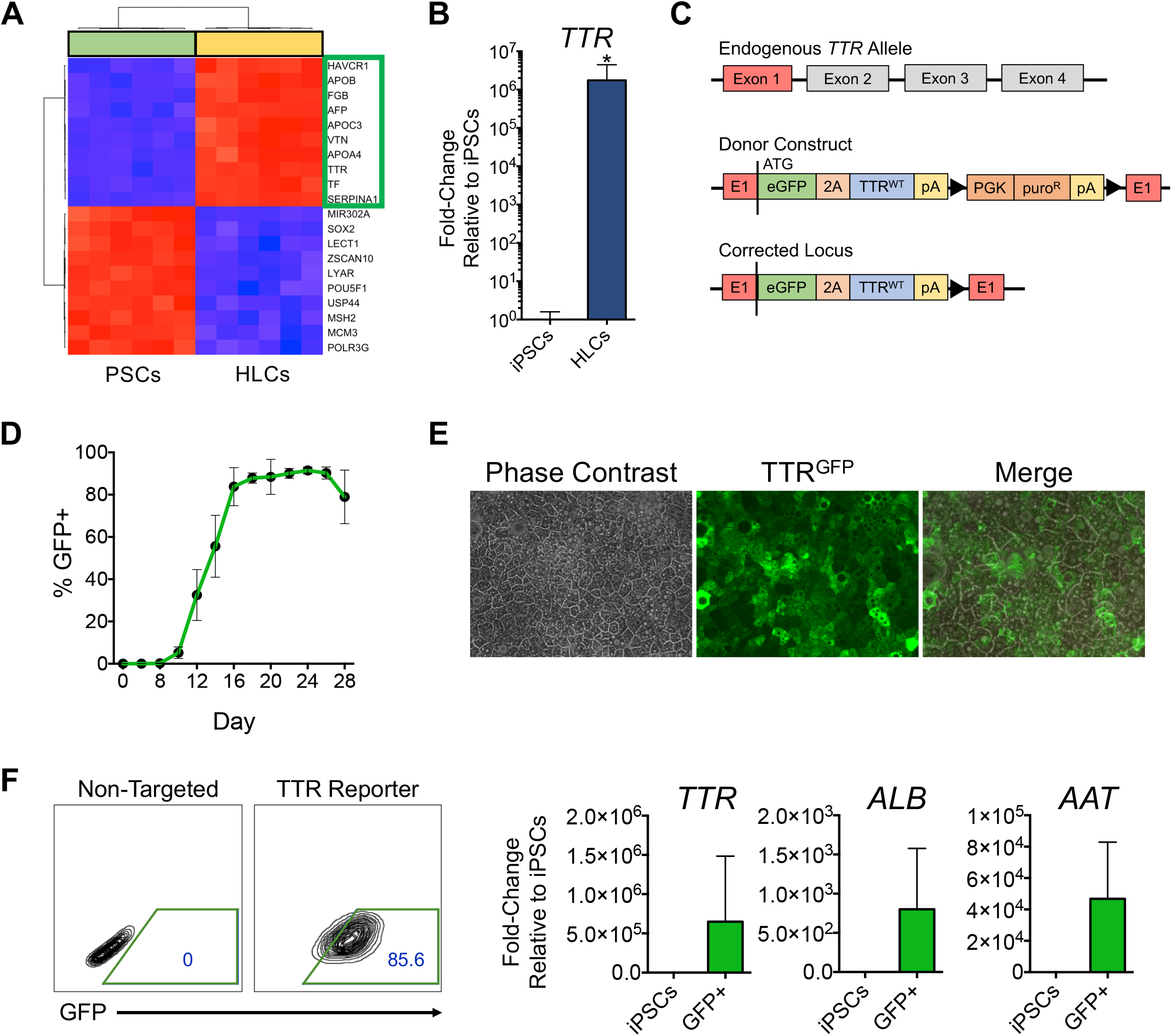
Creation of a TTR promoter-driven hepatic specification reporter iPSC line and universal gene editing strategy for hereditary ATTR amyloidosis. (A) Undifferentiated PSCs and day 24 HLCs (green and yellow columns, respectively) form distinct, independent clusters by microarray analysis. Top 10 transcripts upregulated in HLCs are labeled on the y-axis, highlighted by green box. Top 10 transcripts downregulated in HLCs are labeled on the y-axis (below). Top differentially expressed genes were determined by one-way ANOVA. (B) qRT-PCR validating microarray finding that expression of *TTR* mRNA is significantly upregulated in day 24 HLCs compared to undifferentiated iPSCs; fold-change calculated over undifferentiated iPSCs (n=3, *p<0.05, unpaired t-test for significance, bars denote standard deviation). (C) Schematic representation of the gene targeting strategy. Black triangles flank Cre-excisable LoxP sites. (D) Flow cytometry-based time course of GFP^+^ cells that appear throughout hepatic specification of targeted iPSCs (n=3, bars denote standard deviation). (E) Phase (left) and fluorescence (middle, right) microscopy images of day 26 reporter iPSC-derived HLCs. Images taken at 20X magnification. (F) Expression of hepatic markers in sorted day 16 GFP^+^ HLCs; fold-change calculated over undifferentiated iPSCs (n=5, bars denote standard deviation).

### Gene editing of ATTR amyloidosis iPSCs eliminates secretion of mutant TTR^L55P^ and decreases target cell toxicity

As noted above, *TTR* is one of the most differentially expressed genes in HLC differentiations, suggesting it might serve as an excellent candidate locus to target and generate a hepatic specification reporter iPSC line. To this end, we employed TALEN-mediated gene editing to manipulate an iPSC line derived from a patient with the Leu55 → Pro (TTR^L55P^) mutation, the most proteotoxic disease-causing variant (34, 35). In order to implement a broadly applicable gene correction strategy, we targeted the ATG start site of the endogenous, mutant *TTR* allele, introducing a normal TTR coding sequence, followed by a 2A self-cleaving peptide and *eGFP* coding sequence (**Fig. 1C**). Inclusion of a 2A peptide allows for transcription of a single mRNA that ultimately results in two independent TTR and GFP proteins via a post-translational cleavage event. As a result of this targeting methodology, transcription and translation of mutant TTR is abrogated via introduction of an artificial STOP codon and polyA sequence, and replaced with a normal TTR coding sequence (**Fig. 1C**). Importantly, this universal gene correction strategy provides a singular technique for correcting all known *TTR* genetic lesions while simultaneously obviating concerns regarding haploinsufficiency via replacement of the endogenous mutant *TTR* allele with a wild-type copy. (Additional information regarding generation and characterization of corrected iPSCs can be found in **Supplemental Fig. 1**.)

Using this TTR reporter line, we then measured the kinetics of GFP expression throughout HLC differentiation by flow cytometry. In doing so, we found that expression of GFP peaked at approximately day 16 of a 24-day specification protocol (**Fig. 1D**). By day 24 of the differentiation, HLCs exhibited cobblestone-like morphology, and the majority of cells expressed GFP (**Fig. 1E**). To further validate this reporter line and ensure that GFP expression correlated with the expression of TTR as well as other hepatic-specification markers, day 16 GFP^+^ HLCs were sorted and assayed via qRT-PCR. GFP^+^ cells expressed high levels of *TTR*, as well as other hepatic specification markers such as *AAT*and *ALB* (**Fig. 1F**), suggesting that our corrected reporter cell line labels maturing hepatic-lineage cells during specification.

We further examined the ability of this strategy to eliminate the production of destabilized, disease-causing TTR, as well as alleviate downstream toxicity (**outlined in Fig. 2A**). Normal, corrected, and non-targeted, heterozygous TTR^L55P^ iPSCs were differentiated into HLCs. Conditioned supernatant from each line was harvested after culturing cells for 72 hours in hepatic specification media beginning on day 16 of the differentiation. We used liquid chromatography (LC)/mass spectrometry (MS) to show that TTR immunopurified from normal hepatic supernatants contained TTR^WT^, but not TTR^L55P^, while supernatants collected from patient iPSC-derived HLCs contained both TTR^WT^ and TTR^L55P^ (**Fig. 2B**). Supernatant from corrected iPSC-derived HLCs revealed complete elimination of TTR^L55P^, while levels of TTR^WT^ remained unperturbed (**Fig. 2B**). Importantly, the two amino acid overhang on the N-terminal portion of TTR, resulting from the post-translational cleavage of the 2A peptide, was removed with the TTR signal peptide through normal protein processing in the ER, evident by the identical molecular weights observed for endogenous and exogenous TTR^WT^. This shows that TTR^WT^ from our donor construct is indistinguishable from the endogenous protein.

**FIGURE 2.**
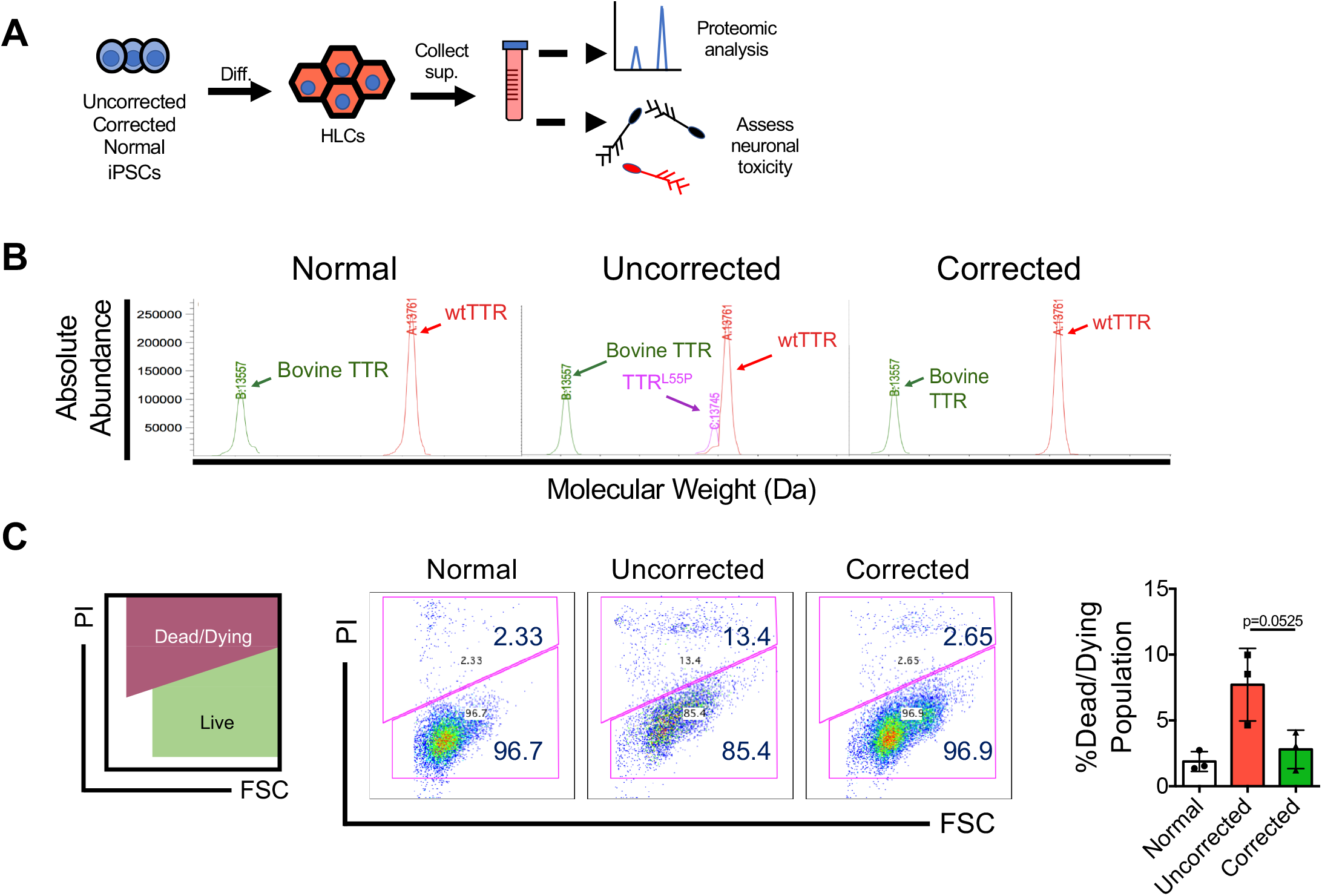
Gene-edited iPSC-derived HLCs no longer produce neurotoxic, destabilized TTR variants. (A) Experimental overview depicting interrogation of normal, TTR^L55P^, and corrected iPSC-derived HLC supernatants and determination of their requisite downstream effects on neuronal target cells. (B) LC/MS analyses of supernatant from normal, TTR^L55P^, and corrected iPSC-derived HLCs. Red trace: TTR^WT^, pink trace: destabilized TTR^L55P^ variant. Bovine TTR (green) is present in media supplements. The molecular weight of each species is denoted in daltons. (C) SH-SY5Y cells were dosed for 7 days with conditioned iPSC HLC-derived supernatant from normal, TTR^L55P^, or corrected conditions. Cell viability was determined via PI staining (n=3, unpaired t-test for significance comparing uncorrected and corrected conditions, bars denote standard deviation).

Many studies have shown that decreasing circulating levels of destabilized TTR species, as in the case of liver transplantation and our novel gene editing strategy, results in decreased peripheral organ dysfunction (36–38). Therefore, we sought to determine the efficacy of our iPSC-based gene correction methodology in decreasing toxicity in a cell-based model. To accomplish this, a neuroblastoma cell line (SH-SY5Y) was dosed with conditioned supernatant generated from mutant TTR^L55P^, corrected, or normal HLCs, and surveyed for toxicity. In these assays, SH-SY5Y cells dosed with mutant hepatic supernatant displayed an increase in PI^+^ cells compared to those dosed with normal supernatant (**Fig. 2C**). Cells dosed with corrected supernatant, however, exhibited a decrease in toxicity comparable to levels observed in the normal control sample (**Fig. 2C**). These results suggest that the proposed gene correction strategy ameliorates TTR-mediated toxicity via reductions in the hepatic secretion of destabilized TTR.

### Single cell RNA sequencing reveals a novel hepatic gene signature associated with the production of destabilized TTR^L55P^

Historically, it has been thought that the livers of patients with ATTR amyloidosis are unaffected during disease pathogenesis (1–9). Recent work however, calling into question the use of donor organs from ATTR amyloidosis patients for DLT procedures challenges this notion (21–26). Furthermore, evidence highlights a significant potential role for the liver in regulating the extracellular aggregation and distal deposition of TTR implicated in ATTR amyloidosis disease pathogenesis (27). Collectively, these results indicate that genetic or aging-related perturbations to the liver could influence the toxic extracellular aggregation and deposition of TTR aggregates on peripheral target tissues.

In order to define specific hepatic proteins and pathways associated with the production of destabilized, amyloidogenic TTR variants, we coupled our TTR reporter system with single cell RNA sequencing (scRNAseq) to compare mRNA expression profiles in syngeneic iPSC-derived HLCs with or without the TTR^L55P^ mutation. In addition to our corrected TTR reporter iPSC line, we also constructed a reporter cell line where our TTR-GFP donor construct was targeted to the wild-type TTR allele in the same TTR^L55P^ parental iPSC line. As a result, we created two syngeneic, TTR-promoter driven hepatic-specification reporter iPSC lines, where the only difference is the presence or absence of the disease causing TTR^L55P^ mutation (henceforth referred to as uncorrected and corrected cells, respectively). To compare HLCs +/− TTR^L55P^, uncorrected and corrected reporter iPSCs were subjected to our hepatic specification protocol until TTR expression had plateaued (at day 16 of the differentiation) (**Fig. 1D**). To control for the inherent heterogeneity of iPSC differentiations, GFP^+^ cells were purified by FACS to select for cells undergoing hepatic-specification (i.e. at similar stages in their developmental trajectories). Transcriptomic profiling was subsequently performed at single cell resolution via the Fluidigm C1 platform (**outlined in Fig. 3A**).

**FIGURE 3.**
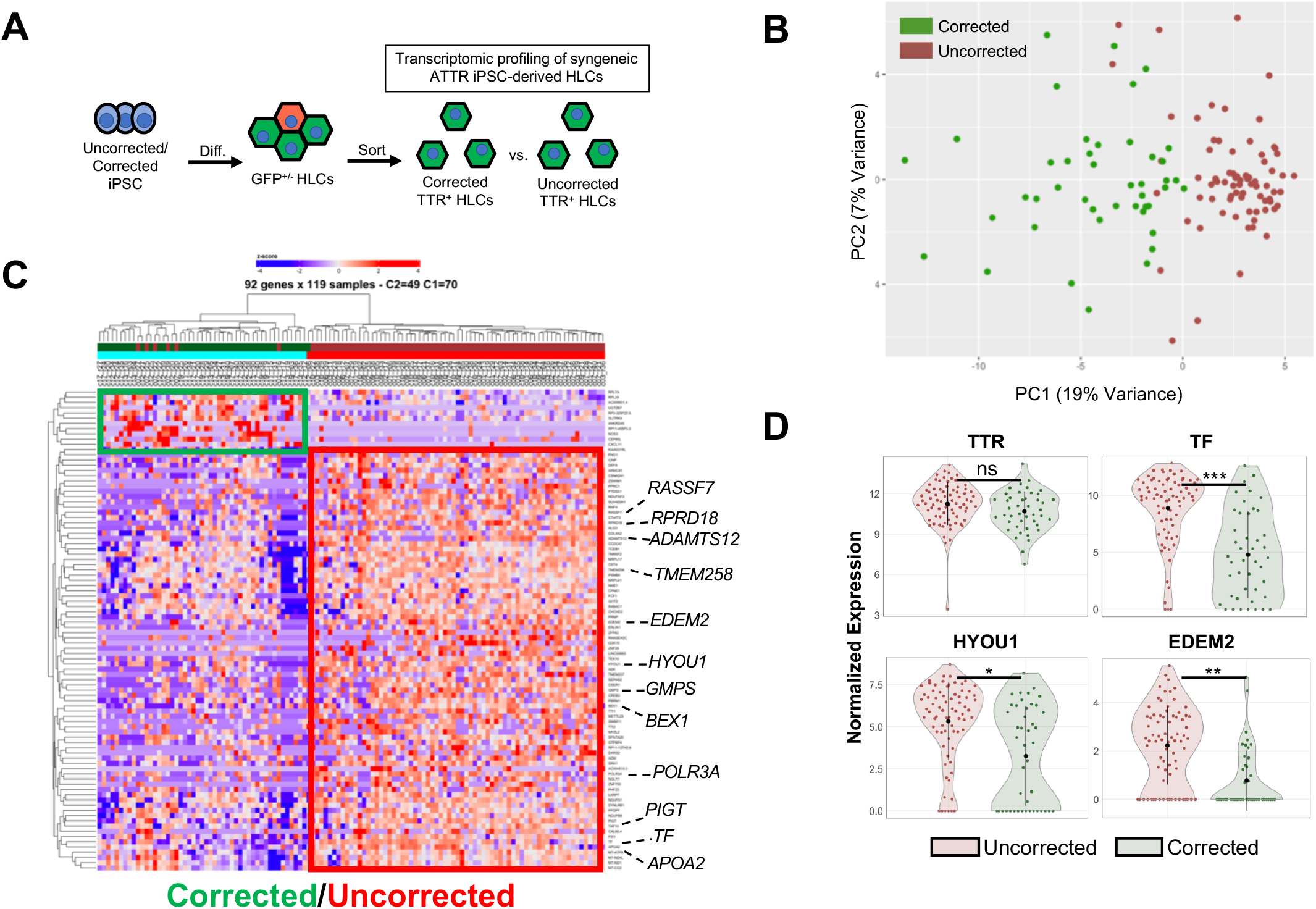
Single cell RNA sequencing (scRNAseq) of corrected vs. uncorrected syngeneic iPSC-derived HLCs reveals a novel hepatic gene signature. (A) Experimental schematic for the transcriptomic comparison of uncorrected (TTR^L55P^-expressing) and corrected syngeneic iPSC-derived HLCs at day 16 of the hepatic specification protocol. (B) Uncorrected (red) and corrected (green) populations form distinct groups by supervised principal component analysis (PCA). Supervised PCA was constructed using the top 500 differentially expressed genes by FDR. (C) Heatmap depicting the 92 genes differentially expressed between uncorrected and corrected populations (one-way ANOVA, FDR cutoff<0.05). Columns represent individual cells, green bar denotes corrected cells, red bar denotes uncorrected cells. Rows represent differentially expressed genes. The top 10 genes by foldchange (uncorrected over corrected) as well as proteostasis factor *EDEM2* are highlighted on the y-axis. (D) Violin plots representing relative expression levels of *TTR*, potential mediators of TTR fibrillogenesis (*TF*), and UPR target genes (*HYOU1, EDEM2*). (FDR determined via one-way ANOVA, *FDR<0.05, **FDR<0.005, ***FDR<0.0005.)

Day 16 uncorrected and corrected HLCs formed clear and distinct groups by supervised principal component analysis (PCA), with 92 genes differentially expressed between the two groups (**Fig. 3B–3D, Supplemental Data File 1**) (significance determined via one-way ANOVA, FDR cutoff <0.05). These analyses identified increased expression of distinct genes and pathways previously shown to influence extracellular aggregation of destabilized TTRs in uncorrected but not corrected HLCs (vide infra) (28–32).

### Transferrin expression is significantly increased in uncorrected HLCs and may represent a novel chaperone for destabilized TTR

The top differentially expressed gene in uncorrected HLCs is the iron transporter, transferrin (TF) (**Fig. 3C, 3D**). Although TF is a known hepatic lineage marker, no other hepatic markers (including: *TTR, ALB, AFP, HNF4A, FOXA1, GATA4, SERPINA1, FGB, DUOX2, A2M, TGM2, HAVCR1, GATA6*) are differentially expressed between corrected and uncorrected HLCs, suggesting that the differential expression of *TF* is not simply due to the differentiation status of individual lines (**Fig. 3D** and **Supplemental Fig. 2**).

Interestingly, previous studies have demonstrated the ability of TF to act as a chaperone in the context of other amyloid disorders such as Alzheimer’s Disease (AD) as well as to physically interact with TTR fibrils *in vivo* (39). As a result, we sought to assess the capacity for TF to act as a novel chaperone for misfolding TTR and in turn prevent TTR fibril formation. To this end, we performed an *in vitro* fibril formation assay whereby the formation of congophilic fibrils from bacterial-derived, recombinant TTR^L55P^ was assessed with or without the addition of TF (**Fig. 4A**). In doing so, iron-free TF (apo-transferrin) at physiologically-relevant concentrations (30 μM) reduced the amount of congophilic species formed by approximately 60% (**Fig. 4B**). This suggests that the increased expression of TF in HLCs containing mutant TTR^L55P^ could represent a mechanism to suppress aggregation-associated toxicity.

**FIGURE 4.**
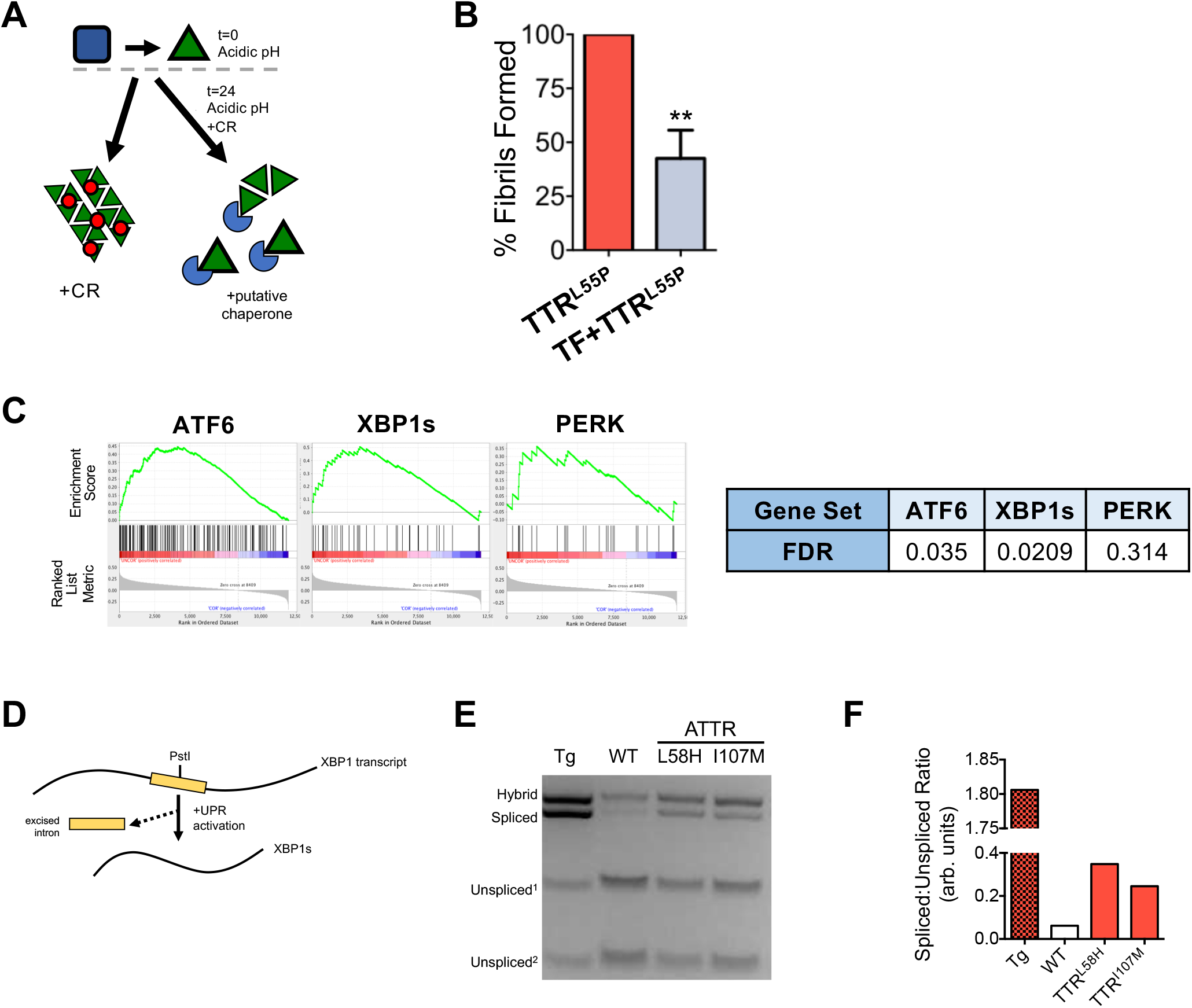
Assessment of TF chaperone capacity and functional validation of XBP1 activation in ATTR amyloidosis hepatic cells. (A) Experimental outline for assessing TF’s *in vitro* ability to prevent the formation of congophilic species from recombinant TTR^L55P^. (B) Percentage of TTR^L55P^ fibrils formed as determined by amount of Congo red (CR) bound after 24-hour incubation of recombinant protein under fibril forming conditions (n=5, **p<0.005, unpaired t-test for significance comparing apo-TF condition to TTR^L55P^ alone). (C) GSEA depicting significant enrichment of adaptive UPR machinery (ATF6, XBP1s) but not PERK target genes in uncorrected HLCs. In these analyses, 100 uncorrected and 60 corrected cells were studied. (D) Depiction of XBP1 splicing in the presence of ER stress and UPR activation. (E) PstI analytical digest of amplified XBP1 transcripts from iPSCs treated with thapsigargin (Tg), wild-type iPSC-derived HLCs (WT), and ATTR HLCs differentiated from two patient-specific iPSC lines (L58H and I107M). Hybrid band represents a PstI-resistant spliced-unspliced XBP1 product generated via PCR protocol. (F) Densitometric quantitation of PstI-digested XBP1 transcripts. Ratio determined by 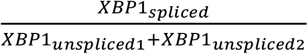.

### Uncorrected HLCs show increased activation of protective UPR-associated signaling pathways

Apart from *TF*, our scRNAseq analysis also identified increased expression of multiple UPR-regulated ER proteostasis factors (e.g., *HYOU1* and *EDEM2;* **Fig. 3C, 3D**) in iPSC-derived HLCs expressing TTR^L55P^. These genes play important roles in regulating proteostasis within the ER. Thus, the increased expression of these ER proteostasis factors suggests that the presence of the destabilized TTR^L55P^ protein challenges the ER proteostasis environment and in turn activates the UPR.

Interestingly, ER stress and UPR activation can influence the toxic extracellular aggregation of amyloidogenic TTR mutants implicated in ATTR amyloidosis disease pathogenesis (28–32). Chemical toxins that induce severe, unresolvable ER stress in mammalian cells decrease the population of amyloidogenic TTR secreted as the native TTR tetramer and instead increase TTR secretion in non-native conformations that rapidly aggregate into soluble oligomers implicated in ATTR amyloidosis disease pathogenesis (28, 40). In contrast, enhancing ER proteostasis through the stress-independent activation of the adaptive UPR-associated transcription factor ATF6, selectively reduces the secretion and subsequent aggregation of destabilized, amyloidogenic TTR variants (29–32). This suggests that activation of adaptive UPR signaling pathways independent of severe ER stress is a protective mechanism to suppress the secretion and toxic aggregation of destabilized, amyloidogenic TTR mutants.

In order to better define the impact of TTR^L55P^ expression on ER stress and UPR activation, we used gene set enrichment analysis (GSEA) to define the extent of UPR pathway activation in our uncorrected iPSC-derived hepatic lineages. Notably, this analysis revealed modest activation of the adaptive IRE1/XBP1s and ATF6 UPR transcriptional signaling pathways, with no significant activation of the pro-apoptotic PERK UPR pathway (**Fig. 4C**). We confirmed IRE1/XBP1s activation in two independent patient-specific iPSC-derived HLCs by monitoring IRE1-dependent *XBP1* splicing (**Fig. 4D-F**). As a positive control, cells were dosed with global UPR activator thapsigargin. These results show that expression of destabilized, amyloidogenic TTR^L55P^ promotes adaptive remodeling of ER proteostasis through the IRE1/XBP1s and ATF6 UPR signaling pathways.

### Hepatic activation of ATF6 signaling selectively reduces secretion of destabilized TTR^L55P^

We next determined the consequence of functional activation of adaptive UPR-associated signaling pathways in ATTR amyloidosis patient-specific iPSC-derived hepatic cells expressing mutant, destabilized TTR. To accomplish this, we introduced an ATF6-inducible donor construct into our previously described heterozygous TTR^L55P^ patient-specific iPSC line. In these cells, the coding sequence for the active N-terminal bZIP transcription factor domain of ATF6 is fused to a destabilized dihydrofolate reductase (DHFR) tag as previously described (32). In the absence of chemical chaperone trimethoprim (TMP), the DHFR.ATF6 protein product is targeted for degradation via the ubiquitin proteasome system (**Fig. 5A**). Upon administration of TMP, the DHFR domain is stabilized, allowing dosable, stress-independent activation of ATF6 transcriptional activity (**Fig. 5A**). ATF6-inducible iPSCs were differentiated into HLCs and subsequently dosed with TMP, beginning on day 15 of hepatic specification. Administration of TMP induced selective expression of the ATF6 target genes *HSPA5* and *HERPUD1*, but not an IRE1/XBP1s target gene (e.g., *ERDJ4*) or PERK target gene (e.g., *GADD34*) (**Fig. 5C, 5D**), confirming selective TMP-dependent ATF6 activation in these HLCs. We then collected conditioned media for 72 hours on patient iPSC-derived HLCs dosed with or without TMP and monitored the relative populations of TTR^WT^ and TTR^L55P^ by mass spectrometry. We initially showed that the relative recovery of TTR^L55P^ from IPs of media prepared on HLCs treated with TMP was reduced relative to TTR^WT^, suggesting reduced secretion of this destabilized TTR variant induced by stress-independent ATF6 activation (**supplemental Fig. 3A, 3B**). To better quantify this reduction, we performed Tandem Mass Tag (TMT) / LC-MS/MS quantitative proteomics to directly monitor the relative amounts of peptides comprising TTR^WT^ or TTR^L55P^ in these conditioned media (**Fig. 5E**). Using this quantitative approach, we show that TMP-dependent ATF6 activation preferentially reduces levels of destabilized TTR^L55P^ 25% relative to TTR^WT^ in HLC conditioned media. This demonstrates that stress-independent ATF6 activation selectively reduces secretion of destabilized, amyloidogenic TTR^L55P^ in patient iPSC-derived HLCs. We moreover found that branch-specific activation of ATF6 signaling protects iPSC-derived HLCs from morphological defects resulting from prolonged exposure to severe ER stress via addition of thapsigargin (**supplemental Fig. 4A, 4B**).

**FIGURE 5.**
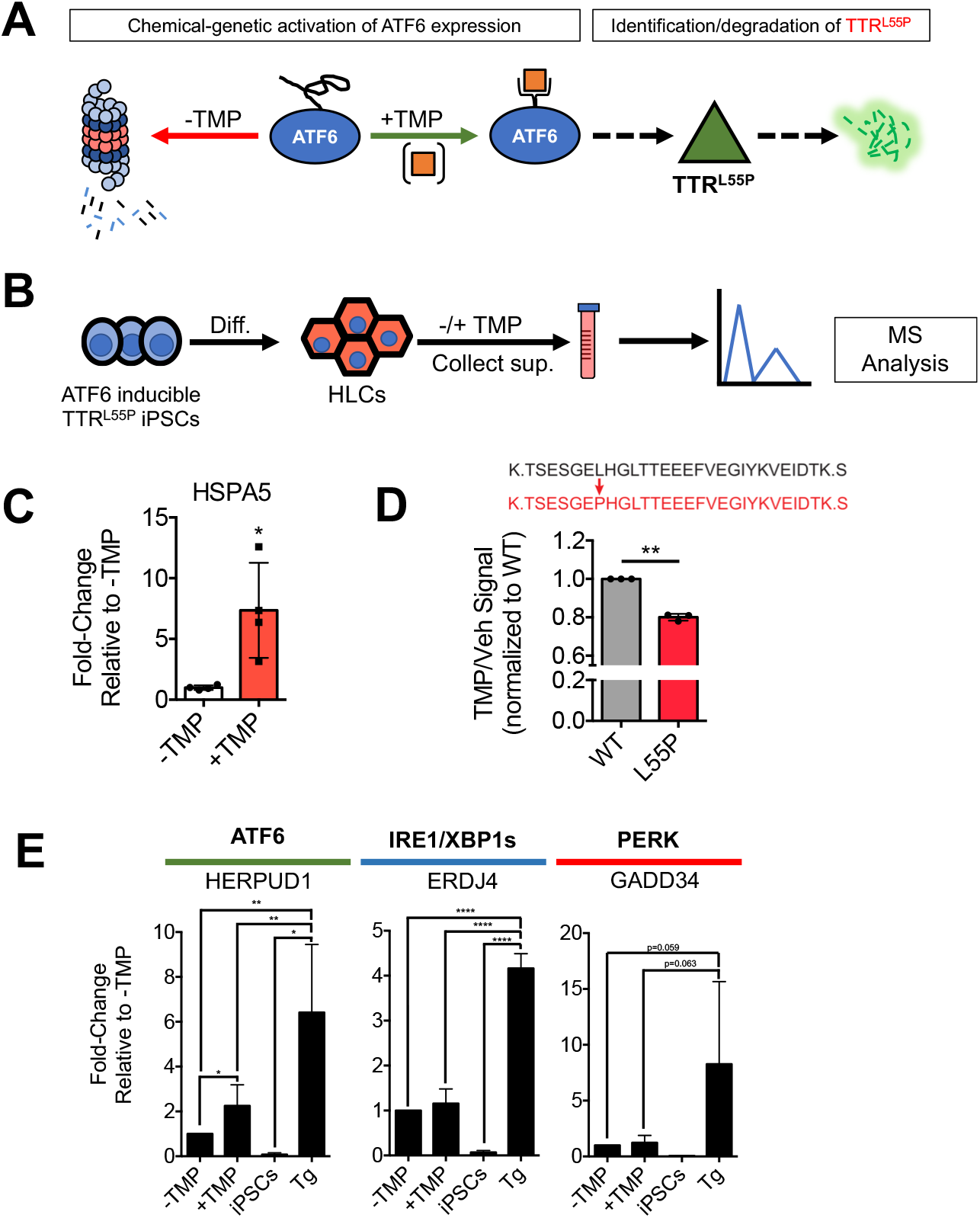
Hepatic stress-independent, branch-specific activation of adaptive UPR-associated ATF6 signaling results in the targeting and selective reduction in the secretion of destabilized TTR^L55P^. (A) A chemical inducible system for activating ATF6 signaling in TTR^L55P^ iPSC-derived cell types. In the absence of chemical chaperone TMP, DHFR.ATF6 is degraded. Upon addition of TMP, DHFR.ATF6 is stabilized and targets and attenuates the secretion of misfolded TTRs. (B) ATF6-inducible iPSCs were differentiated into HLCs. TMP was added and conditioned supernatant was collected and interrogating for the presence and relative abundance of different TTR species via LC/MS-MS. (C) ATF6 target gene *HSPA5* was found to be significantly upregulated upon addition of TMP by qRT-PCR (n=4, *p<0.05, unpaired t-test for significance comparing -TMP to +TMP conditions). (D) LC-MS/MS was used to directly detect the presence of TTR^WT^ (upper peptide sequence, black) and TTR^L55P^ (lower peptide sequence, orange) peptides in conditioned supernatant in the presence and absence of TMP. Abundance of TTR^L55P^ was found to significantly decrease upon activation of ATF6 signaling by ~25% relative to TTR^WT^. Quantities of each peptide were normalized to TTR^WT^ (n=3, **p<0.05, unpaired t-test for significance comparing normalized quantities of TTR^WT^ and TTR^L55P^). (E) Upon addition of TMP, ATF6 target gene *HERPUD1* was found to be significantly upregulated compared to DHFR.ATF6 HLCs in the absence of TMP. IRE1/XBP1s and PERK target genes *ERDJ4* and *GADD34* however, were not found to be differentially expressed in the presence of TMP. Positive control thapsigargin (Tg) was found to significantly upregulate expression of all UPR target genes tested (n=6, *p<0.05, **p<0.005, ****p<0.0001).

## DISCUSSION

Herein, we describe the utilization of gene editing technology in ATTR amyloidosis patient-specific iPSCs in order to uncover a novel human hepatic gene signature resulting from the expression of destabilized, misfolding-prone TTR. Historically, the livers of patients with ATTR amyloidosis were thought to be unaffected in disease pathogenesis (1–6). This is best highlighted through the employment of DLTs, wherein the livers of patients with ATTR amyloidosis are removed and given to individuals in end-stage liver failure. Despite routine use of these transplants, recent clinical data demonstrates that non-ATTR amyloidosis DLT recipients not only go on to develop amyloidogenic TTR fibrils (and disease), but they do so at a median time of only 7.5 years post-transplant (21–26). While these clinical observations suggest a role for the liver in contributing to pathogenesis of ATTR amyloidosis and potentially other systemic amyloid diseases, the cellular and molecular mechanisms for this phenomenon remain unknown.

Through the use of gene editing and scRNAseq, we defined distinct transcriptional profiles for hepatic cells expressing destabilized TTR^L55P^. We hypothesized that hepatic production of destabilized TTRs results in the upregulation of stress-responsive proteostasis factors that regulate the secretion and subsequent aggregation of destabilized TTR variants such as TTR^L55P^. Our scRNAseq experiment revealed that uncorrected hepatic cells exhibited differential expression of 92 genes compared to syngeneic hepatic cells where the only difference is the absence of expression of the mutant TTR. In line with our rationale, we identified many instances in which mutant hepatic cells upregulated expression of well-documented ER stress-associated proteostasis factors involved in regulating protein secretion (e.g., *HYOU1, EDEM2)*.

Interestingly, *TF* was found to be the most upregulated gene in uncorrected, TTR^L55P^-expressing hepatic cells. Despite limited prior evidence for TF as a chaperone for misfolded TTRs, recent work implicates its chaperone capacity in other amyloid disorders such as Alzheimer’s Disease (AD), noting increased protein-level expression in the prefrontal cortices of AD patients compared to elderly, non-diseased individuals (41). Moreover, recent work demonstrated the ability of TF to physically interact with and prevent the self-assembly and toxicity of amyloid ß peptide (Aß) oligomers, the amyloidogenic protein species in AD (42, 43). At the same time, recent *in vivo* data have demonstrated physical interactions between TF and TTR amyloid fibrils (39). Through the use of congophilic fibril formation assays, we demonstrated that iron-free TF at physiologically relevant levels, decreased *in vitro* TTR^L55P^ fibril formation by approximately 60%, implicating TF as a novel chaperone for hepatic TTRs. These observations, together with our scRNAseq and biochemical data, suggest the possibility that TF plays a similar protective role in ATTR amyloidosis. In this model, hepatic cells producing mutant TTR may express higher levels of TF to prevent toxicity and/or fibril formation.

In addition to the differential expression of known and novel chaperone genes, we noted activation of the adaptive arms of the UPR (ATF6 and IRE1/XBP1s) in hepatic cells expressing mutant TTR. Together, these data indicate that the expression of TTR^L55P^ in iPSC-derived HLCs does not induce severe ER stress, but suggests that the activation of adaptive IRE1/XBP1s and ATF6 signaling observed in these cells reflects a protective mechanism to suppress secretion and subsequent aggregation of the destabilized TTR^L55P^ protein. Consistent with this, our IP-based LC-MS analysis of conditioned media from uncorrected HLCs showed that TTR^L55P^ levels were approximately 30% that of TTR^WT^ (**Fig. 2B**). This result mirrors the lower levels of destabilized TTR mutants (as compared to wild-type TTR) observed in conditioned media prepared on hepatic cells expressing both variants (18–20, 28–32).

Our scRNAseq experiments demonstrate dynamic activation of proteostasis transcriptional networks consisting of upregulation of chaperones as well as functional activation of adaptive UPR-associated signaling pathways in mutant HLCs. Due to the gradual nature of the deposition of TTR aggregates and fibrils, with disease manifesting clinically around the 5^th^ or 6^th^ decade of life, ATTR amyloidosis is widely considered an aging-related disorder. It is well-understood that proteostasis factors and the ability to cope with the production of misfolded proteins decreases with age, while similarly, iron has been shown to increase in a number of organs throughout aging (41–46). Interestingly, since UPR signaling declines during normal aging (47–50), the presence of adaptive UPR signaling in the above-mentioned HLCs could reflect protective biologic pathways whose activity also decline during the aging process. Aging-dependent reductions in adaptive UPR signaling could exacerbate TTR^L55P^-associated ER stress and increase secretion of TTR in non-native conformations that facilitate toxic extracellular aggregation. Thus, monitoring changes in hepatic UPR activation and/or conformational stability of circulating TTR tetramers could reflect a potential biomarker to monitor progression of TTR amyloid disease pathogenesis (51) – a notoriously difficult disorder to diagnose.

Together, these results indicate that the expression of a destabilized, aggregation-prone protein upregulates proteostasis factors as well as functionally activates adaptive UPR-associated signaling pathways in ATTR amyloidosis patient-specific iPSC-derived hepatic cells. Moreover, we demonstrated that inducible activation of ATF6 signaling in these cells resulted in a ~25% reduction of destabilized TTR^L55P^ while wild-type levels appeared unaffected. Despite a relatively modest decrease, this reduction could represent a shift toward stability where properly folded, mutant TTRs form stabilized heterotetramers with wild-type TTRs (52).

Conventional therapeutics for many systemic amyloid diseases involve decreasing circulating levels of the amyloidogenic protein. In the case of ATTR amyloidosis, for example, liver transplantation relies upon eliminating circulating levels of mutant TTR. Similarly, emerging RNAi-based therapeutics are being developed to target and eliminate wild-type and mutant TTR transcripts within liver tissue (53–56). Activating ATF6 signaling in ATTR amyloidosis hepatic cells could result in a therapeutic decrease in the secretion of misfolding-prone TTRs from the liver and thus a decrease in extracellular deposition of proteotoxic aggregates as distal target tissues. Recent studies have demonstrated that stress-independent, selective activation of ATF6 signaling is achievable via addition of small molecules (29, 32, 57). At the same time, upregulation of ATF6 signaling is relatively tolerated in humans with hyperactivating mutations (38, 39, 57). Future work will aim to further the development of small molecule-based ATF6-modulating compounds for the treatment of systemic amyloid diseases such as ATTR amyloidosis.

Results from these experiments challenge the long-held notion that the livers of patients with ATTR amyloidosis are unaffected by the disease and do not contribute to pathogenesis. Through the use of our novel iPSC-based model, our work demonstrates that expression of amyloidogenic TTR results in transcriptional and functional changes in ATTR amyloidosis hepatic cells. Moreover, these data suggest that the liver employs protective mechanisms via adaptive UPR-associated signaling pathways in order to cope with the production of misfolding-prone TTRs. Furthermore, this work demonstrates that modulation of UPR-associated ATF6 signaling results in a selective decrease in the secretion of misfolded proteins in patient-specific iPSC-derived hepatic cells, and could potentially represent a broadly applicable therapeutic strategy for complex and diverse systemic amyloid disorders.

## MATERIALS AND METHODS

### Patient samples

To capture the large phenotypic diversity of patients with ATTR amyloidosis, samples were procured from the Amyloidosis Center of Boston University School of Medicine. Reprogramming of material was performed on fresh samples immediately following collection or using frozen mononuclear cells that were previously collected and isolated from subjects.

### iPSC generation and maintenance

Derivation of our patient-specific iPSCs was performed as described (58–60). For all studies described here, cells were maintained either on inactivated mouse embryonic fibroblast (MEF) feeders with KSR supplemented media or under feeder-free conditions using mTeSR1 media. These studies were approved by the Institutional Review Board of Boston University.

### Post-hoc microarray analysis

The heatmap and differential expression analysis was performed using data from Wilson et al. 2015. Differential expression between T24 and T0 of the hepatic differentiation was performed, resulting in 12,129 differentially expressed genes (FDR<0.05). The heatmap was generated using Ward’s hierarchical clustering method on the row-scaled expression values of these genes.

### TALEN-mediated gene editing of patient-specific iPSCs

Cells were transfected via Lipofectamine with PLUS reagent (ThermoFisher, Cat. Nos. 11668019, 11514015). Briefly, iPSCs were cultured until ~60% confluent in a 6-well plate. 1.2 μg of left and right TALEN targeting vectors and 3 μg of donor vector were added to cells. 700 ng/μL puromycin selection was then performed. Puromycin cassette excision was accomplished using transient transfection of the pHAGE-EF1α-Cre-IRES-Neo plasmid followed by subsequent screening and single cell selection and expansion.

### Directed differentiation of iPSCs to hepatocyte-like cells

IPSCs were specified to the hepatic lineage via a 2D feeder-free, chemically-defined differentiation protocol as previously described (18–20, 61, 62).

### RNA isolation, cDNA synthesis, and qRT-PCR

RNA was extracted from cells via RNeasy Mini Kit (Qiagen, Cat. No. 74104). RNA was eluted in RNAse-free H2O and further purified via treatment with DNA removal kit (ThermoFisher, Cat. No. AM1906). ~1 μg purified RNA was used to generate cDNA via High-Capacity cDNA Reverse Transcription Kit (Applied Biosystems, Cat. No. 43-688-13). qRT-PCR was performed with TaqMan Universal Master Mix II, with UNG (ThermoFisher, Cat. No. 4440038). TaqMan Gene Expression Assays used include the following: ß-actin (Hs99999903_m1), AAT (Hs00165475_m1), ALB (Hs00609411_m1), ERDJ4 (Hs01052402_m1), HERPUD1 (Hs01124269_m1), HSPA5 (Hs00607129_gH), GADD34 (Hs00169585_m1), and TTR (Hs00174914_m1). Quantities of genes of interest were compared relative to ß-actin levels. Fold-change was calculated via ΔΔC_⊤_ method. Undetermined C_⊤_ values were taken to be 40. Samples were run in technical duplicate.

### IP-MS analysis of secreted TTR

TTR was immunoprecipitated from conditioned media prepared on iPSC-derived hepatic lineages, as previously described (20). Briefly, we incubated 5 mL of conditioned media overnight at 4°C with cyanogen bromide-activated sepharose covalently conjugated to a rabbit polyclonal anti-TTR antibody (a kind gift from Jeff Kelly, TSRI). We then washed the beads four times with 10 mM Tris pH 8.0, 140 mM NaCl, 0.05% saponin and one time with 10 mM Tris pH 8.0, 140 mM NaCl. TTR was then purified by incubating the beads with 100 μL of 100 mM triethylamine pH 11.5 for 30 min at 4°C. LC/MS analysis was then performed on an Agilent single quadrupole mass spectrometer, as previously described (20).

### Determining downstream neuronal toxicity in response to iPSC-derived HLC supernatant

Conditioned HLC supernatant was generated by incubating hepatic differentiation media on day 16 HLCs for 72 hours. Supernatant was collected and subsequently concentrated using Centrifugal Filter Units (Millipore Sigma, Cat. No. UFC901024). (After first collection, cells were refed with media for an additional 72 hours to generate a second batch of conditioned supernatant.) Supernatant was first centrifuged at 200 x g for 1 minute at room temperature to remove cell debris. Media was then collected, transferred to filter units, and spun at 2140 x g for 45 minutes at room temperature. Concentrated supernatant was subsequently stored at 4°C until dosing experiment. SH-SY5Y cells were plated at 2×10^5^ cells and subsequently dosed for ~7 days with media composed of SH-SY5Y growth media and conditioned supernatant at a 1:1 ratio. Media was replaced every 48 hours until toxicity assay was performed. After dosing cells, floating and adherent SH-SY5Y cells were collected, stained with PI (BD Biosciences, Cat. No. 556463), and analyzed via flow cytometry.

### Single-cell RNA Sequencing and Analysis

Corrected and uncorrected cells were sorted and entered into the Fluidigm C1 HT workflow, which was used to capture and lyse individual cells, reverse transcribe RNA, and prepare libraries for sequencing (See: Using the C1 High-Throughput IFC to Generate Single-Cell cDNA Libraries for mRNA Sequencing, Cat. No. 100-9886). Sequencing was performed on a Nextseq 500 using a high-output kit. This resulted in a total of 430 million paired-end, 75 bp reads. Reads were aligned to the human genome (GRCh38) and quantified using the STAR aligner (63). Outlier removal was performed (cells must have >1,500 genes detected, <3 median absolute deviations away from median total reads, mitochondrial counts), in addition to removing a subpopulation determined through k-means clustering with high mitochondrial expression and low numbers of genes expressed. This resulted in 120 cells available for analysis, with a mean of 482,205 aligned reads/cell. Size factors were computed via the Scran Bioconductor package, which uses pool-based scaling factors and deconvolution, followed by log-transformation normalization using the Scater package (64). A one-way ANOVA was used to determine significantly (FDR <0.05) differentially expressed genes between corrected and uncorrected cells. Supervised PCA was performed using differentially expressed genes as factors. The heatmap was generated using Ward’s hierarchical clustering method (65) on the row-scaled expression values of differentially-expressed genes determined through the aforementioned method. Raw single-cell sequencing data can be accessed from the Gene Expression Omnibus at (GSE number pending).

### XBP1 splicing assay

RNA was isolated from HLCs and cDNA was generated via standard RT reaction (see above). XBP1 transcript was amplified via PCR reaction with forward primer 5’-A AAC AGA GTA GCA GCT CAG ACT GC-3’ and reverse primer 5’-TC CTT CTG GGT AGA CCT CTG GGA G-3’. PCR program utilized included the following steps: 94°C for 4 minutes, 35 cycles of 94°C (10 seconds), 63°C (30 seconds), and 72°C (30 seconds), and lastly 72°C for 10 minutes. Amplified transcripts were subsequently digested with PstI enzyme (New England BioLabs, Cat. No. R0140S) and analyzed on a 2.5% agarose gel. Relative quantities of bands were determined via ImageQuant TL software. For positive control for XBP1 activation, thapsigargin (Millipore Sigma, Cat. No. T9033) was added to undifferentiated iPSCs at a concentration of 1 μM for 24 hours.

### Recombinant expression and purification of TTR^L55P^

TTR^L55P^ was expressed and purified as previously described (66). Briefly, TTR^L55P^ was expressed in *E. coli* as an N-terminal histidine-tagged protein isolated from cell lysate after passage through Ni-NTA agarose (Qiagen, Cat. No. 30210). Bound TTR^L55P^ was then eluted by competition with imidazole. Lastly, purified TTR^L55P^ was loaded into an Econo-Pac 10DG desalting column (BIO-RAD, Cat. No. 7322010) for rapid buffer exchange to 10 mM sodium phosphate buffer pH 7.8, 100 mM KCl, 1 mM EDTA. The final concentration was adjusted to 0.4 mg/ml. Subsequently, purified TTR^L55P^ was stored at 0°C and used within one week.

### In vitro fibril formation assay

TTR^L55P^ fibril formation was triggered under mild acidic conditions. The amount of fibrils formed was measured by Congo red (CR) binding assay as reported previously (67). TTR^L55P^ and lyophilized human plasma-derived apo-transferrin protein (Apo-TF; R&D Sytems, Cat. No. 3188-AT) resuspended in 10 mM sodium phosphate buffer (pH 7.8, 100 mM KCl, 1 mM EDTA) were both filtered through 0.2 μm membranes prior to their incorporation into the reaction mixtures. Addition of 50 mM sodium acetate buffer pH 4.6, 100 mM KCl lowered the pH of the reaction to 4.9. The final concentrations of TTR^L55P^ and Apo-TF were 0.2 mg/ml and 2500 μg/ml respectively. Fibril formation was carried out at 37°C without agitation in a mastercycler with lid temperature of 80°C to avoid condensation. Reactions were halted after 24 hours with the addition of 1.5 M Hepes pH 8.0. Finally, fibril formation reaction was added to 10 μM CR (Sigma-Aldrich, Cat. No. 573-58-0) working dilution (prepared in 5 mM KH2PO4 pH 7.4, 150 mM NaCl). After 15 minutes at room temperature, absorbance was taken at 477 nm and 540 nm. The amount of CR bound to amyloid fibrils was determined via molar concentration of bound CR= A_540nm_/25295 – A_477nm_/46306. The percentage of fibrils formed was calculated via: 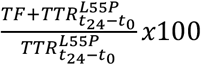.

### Generation of ATF6-inducible TTR^L55P^iPSC line

TTR^L55P^ iPSCs were nucleofected (Lonza) with 3 μg of previously described DHFR.ATF6 donor construct (39) using the manufacturer’s protocol. 48 hours post nucleofection, cells were grown in 500 ng/μL puromycin for approximately 10 days. Successfully grown colonies were subjected to dilution cloning to ensure clonality. To assess functionality, clonal, puromycin-resistant colonies were subjected to 10 μM TMP for 48 hours. RNA was harvested from each clone and qRT-PCR was performed to assess upregulation of ATF6 target gene, *HSPA5*, in the presence of TMP.

### Tandem Mass Tag (TMT)-LC-MS/MS Analysis of TTR^WT^ and TTR^L55P in^ Conditioned Media Prepared on iPSC-derived Hepatic Lineages

Hepatic lineages were prepared from patient iPSCs expressing TTR^WT^ and TTR^L55P^ where TMP-regulated DHFR.ATF6 was introduced. Media was conditioned on these cells for 72 hours in the presence or absence of TMP (10 μM). The media was collected and then subjected to chloroform/MeOH precipitation to precipitate proteins (68). Dried pellets were dissolved in 8 M urea/100 mM TEAB, pH 8.5, reduced with 5 mM tris(2-carboxyethyl) phosphine hydrochloride (TCEP), and alkylated with 50 mM chloroacetamide. Proteins were then dilute to 2 M urea/100 mM TEAB and trypsin digested overnight at 37°C. The digested peptides were labeled with TMT (ThermoFisher, Cat. No. 90309, Lot. No. UB274629). The TMT labeled samples were analyzed on a Orbitrap Fusion Tribrid mass spectrometer (ThermoFisher). Samples were injected directly onto a 30 cm, 100 μm ID column packed with BEH 1.7 μm C18 resin (Waters). Samples were separated at a flow rate of 400 nL/min on a nLC 1200 (ThermoFisher). Buffer A and B were 0.1% formic acid in 5% acetonitrile and 80% acetonitrile, respectively. A gradient of 1–30% B over 160 min, an increase to 90% B over 60 min and held at 90% B for 20 minutes was used for a 240 minute total run time. Column was re-equilibrated with 20 μL of buffer A prior to the injection of sample. Peptides were eluted directly from the tip of the column and nanosprayed directly into the mass spectrometer by application of 2.8 kV voltage at the back of the column. The Orbitrap Fusion was operated in a data dependent mode. Full MS1 scans were collected in the Orbitrap at 120k resolution. The cycle time was set to 3 s, and within this 3 s the most abundant ions per scan were selected for CID MS/MS in the ion trap. MS3 analysis with multinotch isolation (SPS3) was utilized for detection of TMT reporter ions at 30k resolution (69). Monoisotopic precursor selection was enabled and dynamic exclusion was used with exclusion duration of 10 s.

Protein and peptide identification were done with Integrated Proteomics Pipeline – IP2 (Integrated Proteomics Applications). Tandem mass spectra were extracted from raw files using RawConverter (70) and searched with ProLuCID (71) against Uniprot human database. The search space included all fully-tryptic and half-tryptic peptide candidates. Carbamidomethylation on cysteine and TMT labels on N terminus and lysine were considered as static modifications. Data was searched with 50 ppm precursor ion tolerance and 600 ppm fragment ion tolerance. Identified proteins were filtered to 10 ppm precursor ion tolerance using DTASelect (72) and utilizing a target-decoy database search strategy to control the false discovery rate to 1% at the protein level (73). Quantitative analysis was done with Census (74) filtering reporter ions with 20 ppm mass tolerance and 0.6 isobaric purity filter.

Peptide quantifications corresponding to the TSESGE<u>L/P</u>HGLTTEEEFVEGIYKVEIDTK peptide from TTR^WT^ or TTR^L55P^ were initially analyzed for each biological replicate. Peptides showing a m/z signal less than 10,000 were excluded from the analysis. A TMP ratio for each identified peptide was quantified using the following equation: TMP ratio = peptide signal in +TMP channel replicate = n / peptide signal –TMP channel replicate = n. The TMP ratio for individual peptides was then averaged across all peptides observed for each individual biological replicate to generate the plots shown in **Supplemental Fig. 3**. The relative impact of TMP on the secretion of TTR^L55P^ was then quantified by normalizing the average TMP ratio for the TTR^L55P^ peptide to the TMP ratio for the TTR^WT^ peptide to generate the plot shown in **Fig. 5D**.

### Morphological assessment of the impact of ATF6 signaling on HLCs exposed to prolonged ER stress

To assess morphological differences under severe, prolonged stress, iPSC-derived HLCs were dosed with 50 nM thapsigargin for 5 days beginning on day 15 of the differentiation. In stressed cells with ATF6 activation, TMP was simultaneously added at a concentration of 10 μM. In instances where ATF6 signaling was inhibited in the presence of thapsigargin, 6 μM Ceapin-A7 (CP7) (Sigma-Aldrich, Cat. No. SML2330) was added to cells. After dosing for 5 days, cell morphology was noted. To account for degradation, small molecules were supplemented to media at half-concentration every other day.

## STATISTICAL ANALYSIS

Unless otherwise stated in the text, significance was determined between experimental and control conditions via unpaired Student’s t-test. All experiments were performed using technical and biological triplicates. Here, we define biological replicates as individual passages, genetically distinct iPSC lines, and/or separate differentiations. Additional statistical information can be found in individual figure legends.

## AUTHOR CONTRIBUTIONS

RMG designed research studies, conducted experiments, acquired data, analyzed data, and wrote the manuscript. DCL designed research studies, conducted experiments, acquired data, and analyzed data. TMM and NS analyzed data. CTA, JC, SG, JR, JD, and SP conducted experiments. LHC and JRY revised the manuscript. DNK, RLW, and GJM supervised and designed research studies, analyzed data, and revised the manuscript.

## ACKNOWLEDGMENTS

This work was supported by the National Institutes of Health - NIDDK (grant R01DK102635 (GJM, RLW), P41GM103533 (JRY), R01NS092829 (RLW), F31DK121481 (RMG)), NCATS (grant 1UL1TR001430 (RMG, GJM, DNK, AAW), the Amyloidosis Foundation, and the Young Family Amyloid Research Fund.

---

**SUPPLEMENTAL FIGURE 1.**
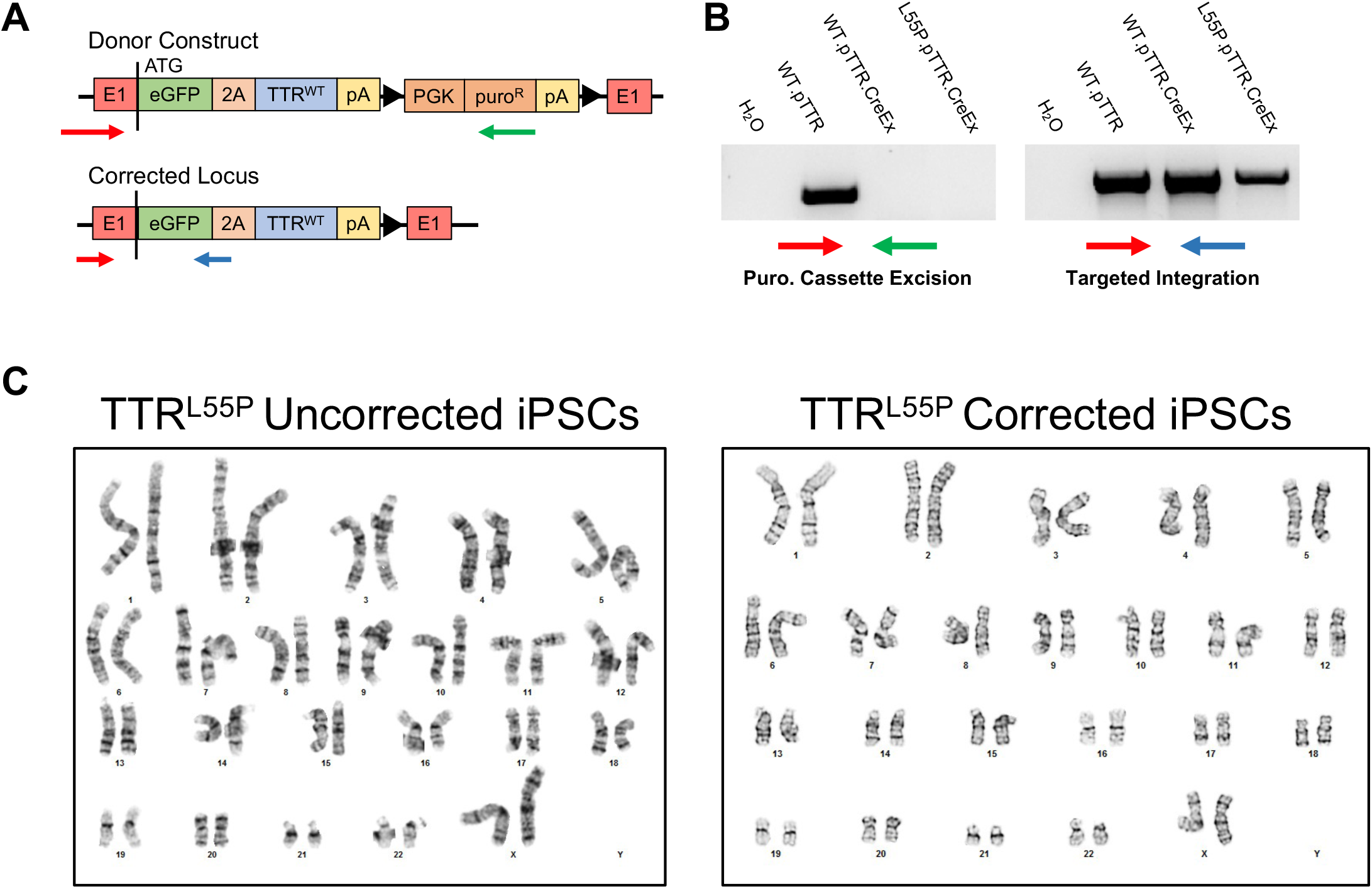
Validation of targeted iPSC reporter line for site-specific integration of transgene as well as karyotypic stability. (A) Targeting construct used to perform gene correction. Red arrows denote the forward primer utilized for allele integration and puromycin cassette excision assay. Blue arrow represents the reverse primer used in the targeted integration screen. Green arrow depicts the reverse primer utilized for puromycin resistance (puro^R^) cassette excision. (B) PCR screen of gDNA for targeted integration of donor construct (left) and excision of puro^R^ cassette (right). (C) Non-targeted (left) and corrected (right) TTR^L55P^ iPSCs derived from a female ATTR amyloidosis patient were karyotypically normal, post-excision of the puro^R^ cassette.

**SUPPLEMENTAL FIGURE 2.**
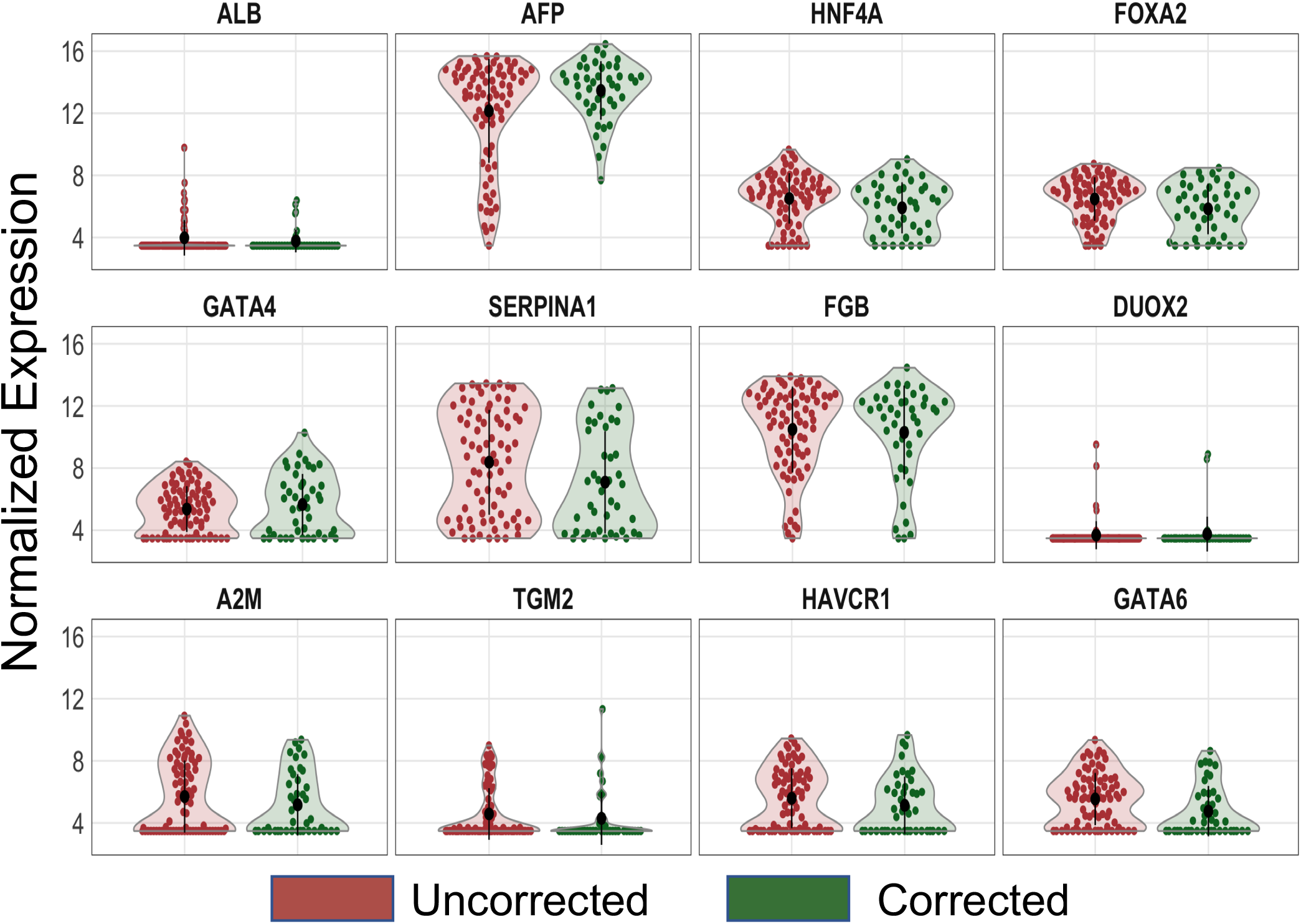
Liver markers are not differentially expressed between uncorrected and corrected HLCs at day 16 of differentiation. Violin plots representing equivalent relative expression levels of genes known to be upregulated during hepatic specification of PSCs in uncorrected (red) and corrected (green) HLCs. None of the noted genes are differentially expressed between the two groups, suggesting competent normalization of the hepatic specification procedure.

**SUPPLEMENTAL FIGURE 3.**
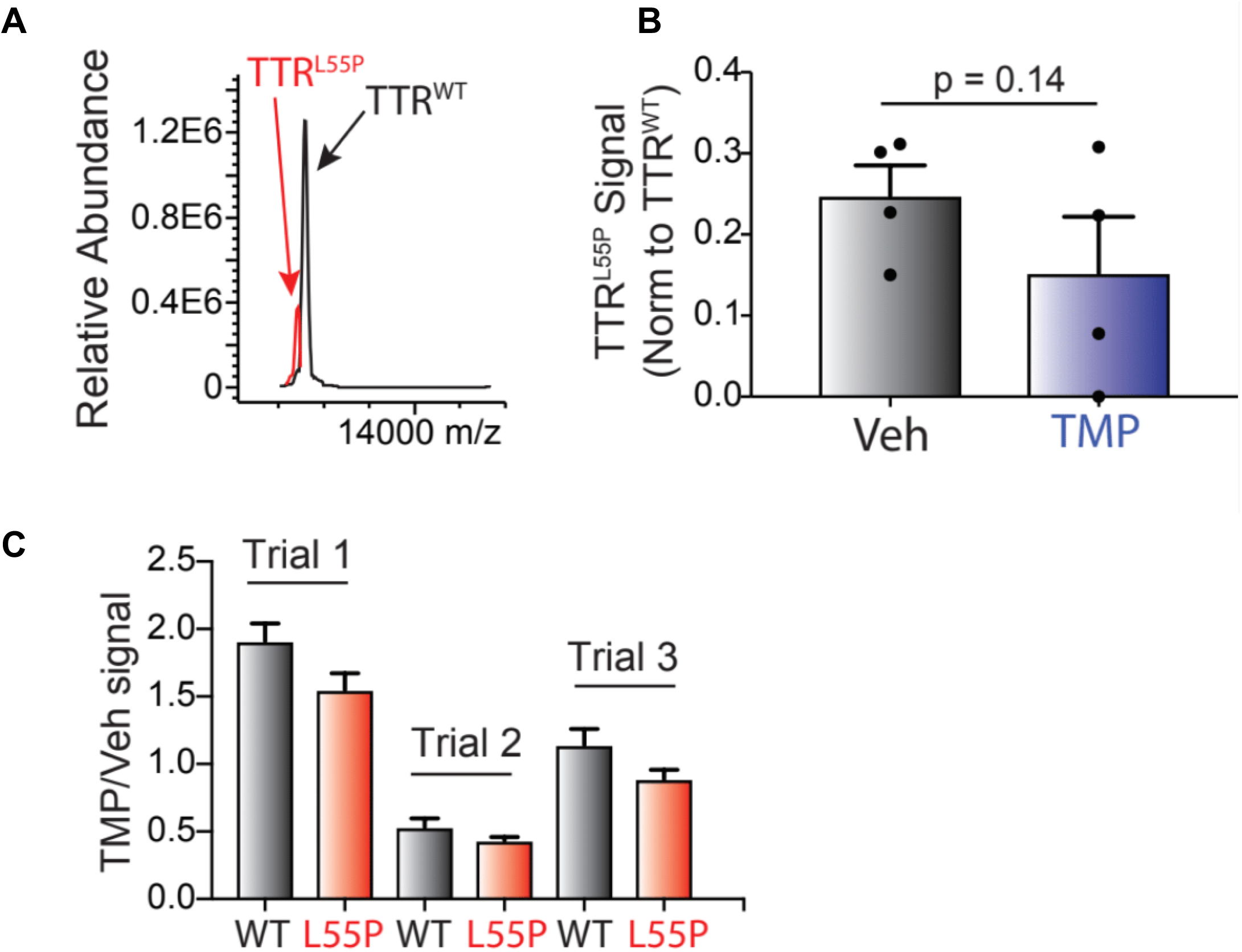
Activation of ATF6 signaling in ATTR amyloidosis patient-specific iPSC-derived HLCs results in selective decrease in the secretion of destabilized TTR. (A) Representative IP-MS data of the relative abundances of TTR^L55P^ and TTR^WT^ in non-targeted, heterozygous TTR^L55P^ patient-specific HLCs. (B) Recovery of TTR^L55P^ in conditioned ATF6-inducible HLC supernatant in the presence and absence of exogenous ATF6 activation (i.e. TMP) (n=4, unpaired t-test for significance). TTR^L55P^ signal denoted is normalized to TTR^WT^ in each replicate. (C) Individual experiments compiled/summarized in **Fig. 5D** of main text. Error bars represent quantification from different spectral counts (2 for TTR^WT^ and 4 for TTR^L55P^) of the same peptide.

**SUPPLEMENTAL FIGURE 4.**
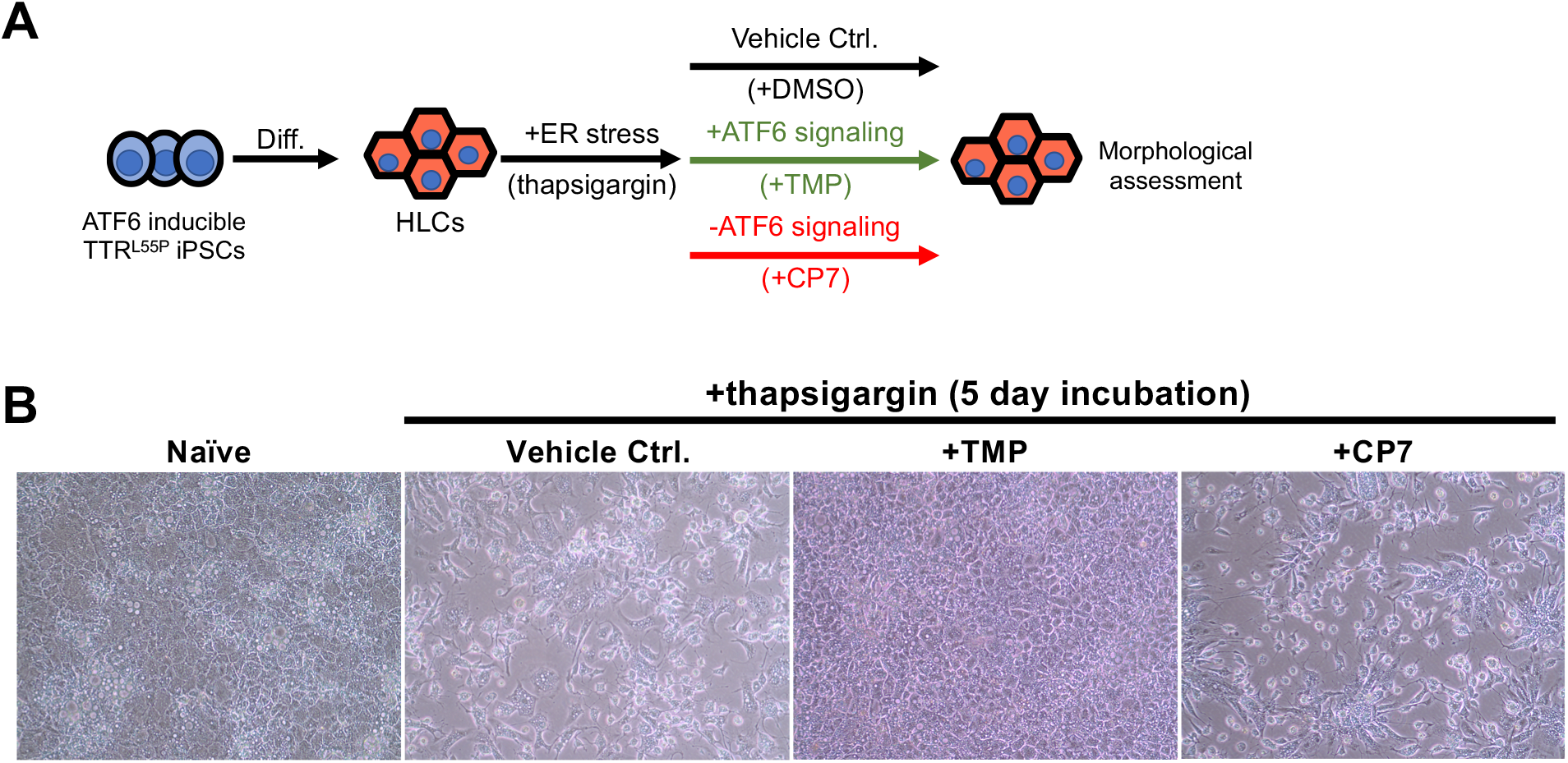
Branch-specific activation of ATF6 signaling in patient-specific iPSC-derived hepatic cells alleviates ER stress-mediated morphological defects. (A) Inducible ATF6 iPSCs were differentiated into HLCs and exposed to global ER stress via addition of thapsigargin. In addition to the presence of thapsigargin, ATF6 signaling was either activated via addition of TMP or inhibited via addition of small molecule CP7. After 5 days of exposure to each compound, cells were observed for morphological defects. (B) In the absence of TMP, in the presence of thapsigargin, cells exhibited gross morphological defects. Upon activating ATF6 signaling (in the presence of thapsgargin), morphology appeared to be rescued. Conversely, upon inhibiting ATF6 signaling via addition of CP7 (in the presence of thapsigargin), morphological defects were noted. Cells were dosed with various compounds in three independent differentiations.

## ABBREVIATIONS

A1AT: alpha-1 antitrypsin
Aß: amyloid ß
AD: Alzheimer’s disease
AFP: alpha fetoprotein
ALB: albumin
ATTR amyloidosis: transthyretin amyloidosis
DLT: domino liver transplant
FDR: false discovery rate
GSEA: gene set enrichment analysis
HLC: hepatocyte-like cell
iPSC: induced pluripotent stem cell
LC/MS: liquid chromatography/mass spectrometry
PCA: principle component analysis
PSC: pluripotent stem cell
scRNAseq: single cell RNA sequencing
TALEN: transcription activator-like effector nuclease
TF: transferrin
TMT: tandem mass tag
TTR: transthyretin
UPR: unfolded protein response

